# Distinct Roles for SETα and SETβ in Early Cell Fate Decisions

**DOI:** 10.1101/2024.09.21.614244

**Authors:** Patrick Siang Lin Lim, Eran Meshorer

## Abstract

SET, the nuclear proto-oncogene, is primarily expressed as SETα in embryonic stem cells. Upon pluripotency exit, a transcriptional switch driven by alternative promoters causes SETβ to largely replace SETα expression. Functional distinctions between the two isoforms have been difficult to ascertain, partly due to the redundancy between SETα and SETβ in their protein structure and activity. In this study, we use embryonic stem cells (ESCs) with inducible SET isoform-specific expression to investigate the differences between both SET isoforms. Time-course RNA-seq analyses in SET-KO backgrounds as well as isoform-specific ChIP-seq experiments reveal regulatory functions for SETα and SETβ. Despite sharing many binding sites and binding partners, SETα has unique regulatory functions on its target genes, while SETβ downregulates FGF4. As KLF5 specifically regulates SETα, this implicates SET isoform switching at the KLF5/FGF signalling axis during primitive endoderm specification. Together, we propose a model of how distinct roles of SETα and SETβ may regulate cell identity in the early blastocyst.

## Introduction

An important feature of embryonic stem cells (ESCs) is the expression of transcription factors (TFs) that cooperate to maintain pluripotency. Together with the core pluripotency network, a variety of other factors, including chromatin remodelers and epigenetic modifiers have also been shown to support an undifferentiated state. Many of these factors are selectively highly expressed in ESCs, demonstrating that a distinctive regulatory program complements the unique chromatin landscape and structure of the pluripotent genome (Klein and Hainer, 2020; Li and Izpisua Belmonte, 2018; Lim and Meshorer, 2020; Pelham-Webb et al., 2020).

In earlier work, we performed screens for potential regulators of pluripotency using an endogenously tagged fluorescent protein library. These experiments led to the detection of SET, also known as TAF-I or I2PP2A, as a protein that is rapidly downregulated upon differentiation of mouse embryonic stem cells (ESCs) (Harikumar et al., 2017). We characterised a transcriptional switch in differentiating pluripotent cells, where expression of the predominant SET isoform expressed in ESCs, SETα, is replaced by expression of SETβ. This conversion is regulated by two alternative promoters, each of which is bound by a distinct subset of TFs; SETα by pluripotency factors, OCT4, SOX2, KLF2 and NANOG, supporting an active state, and SETβ by CAPG, OGT, TET1 and HDAC2, among others, supporting a poised state (Edupuganti et al., 2017). The absence of SET has been shown by us and others in mouse models to be embryonic lethal (Edupuganti et al., 2017; Kon et al., 2019). Previous experiments in *Xenopus* had also implicated SET in neural tube formation (Rashid et al., 2006). In recent work, we and others identified an interaction between SET and p53 in mouse ESCs (Harikumar et al., 2020; Wang et al., 2016). Following depletion of SET, p53 is activated, leading to upregulation of p53 targets and primitive endoderm markers. This has been shown to be caused by SET’s ability, as a subunit of the INHAT complex, to inhibit acetylation of the p53 C-terminus (Kim et al., 2012; Wang et al., 2016). Moreover, we identified a separate interaction between SET and β-catenin in the suppression of canonical Wnt-signalling (Harikumar et al., 2020). Although these findings indicate that SET plays an essential role in early development, the functional basis for the alternative SET promoter switching that occurs upon pluripotency exit remains unclear.

The usage of alternative promoters is a widespread phenomenon in Eukaryotes and can play important functional roles in regulating transcription in different developmental or tissue-specific contexts (Ayoubi and Van De Yen, 1996). For instance, a subset of IGFR promoters are bound and regulated by C/EBP, allowing for specific expression in the liver (van Dijk et al., 1992). Likewise, each of the three retinoic acid receptor (RAR) genes can be expressed as two different isoforms, which differ in their distribution of expression in developing mouse embryos, contributing to the pleiotropic functions of RAR heterodimers (Leid et al., 1992; Mollard et al., 2000).

Considering that differential regulation via alternative promoter usage may indicate functionality, do SETα and SETβ play different roles in ESCs? And if so, would any functional difference be due to additional intrinsic properties of each isoform or the lack thereof? Answering these questions is complicated by the ostensible redundancy between the two isoforms, which share over 91% of their protein sequence. Both SETα and SETβ possess identical C-terminal acidic domains, which facilitate many of their observed functional roles, including histone chaperone function (Muto et al., 2007) and the ability to bind to unacetylated lysine tails (Yao et al., 2023). Furthermore, both isoforms are capable of potent inhibition of protein phosphatase 2A (PP2A) (Li et al., 1996) and are members of the INHAT complex, which inhibits p300 and PCAF-mediated acetylation (Kutney et al., 2004; Seo et al., 2002, 2001).

Nevertheless, despite their similarity in structure and function, there is evidence to suggest that SETα and SETβ do not play completely identical roles. Early experiments noted a higher specific activity in SETβ for stimulating adenovirus DNA replication (Nagata et al., 1995). In vitro disassociation experiments have suggested that SETα’s distinctive N-terminal region may be autoinhibitory and interfere with its own histone chaperone activity (Kajitani et al., 2017). Additionally, several studies suggest that SETα and SETβ possess developmentally relevant functional specificity. In previous work, knockout of SETα and not SETβ in ESCs leads to teratoma formation upon implantation in immune-deficient mice (Edupuganti et al., 2017). Another recent study found that the reprogramming efficiency of mouse embryonic fibroblasts (MEFs) to induced pluripotent stem cells (iPSCs) is compromised following SET knockdown, which occurs in part due to incomplete silencing of retroviral elements. The authors then showed that overexpression of SETα but not SETβ could rescue the silencing of retroviral elements necessary for robust reprogramming (Bui et al., 2019). Despite these reports, a clear understanding of the distinct role(s) that each isoform plays in ESCs is currently lacking, partly, but crucially, due to the lack of N-terminal specific antibodies preventing analysis of isoform-specific genome-wide distributions.

In order to disentangle the isoform-specific contributions of SETα and SETβ, we used a doxycycline (Dox)-inducible SET overexpression system in mouse ESCs. In these cell lines, addition of Dox allows selective exogenous expression of either SETα or SETβ. In addition, we performed CRISPR Knockout (KO) of SET in these SET-overexpressing cell lines to generate a system where isoform-specific addback of SETα or SETβ in a SET-null background could be performed in a controllable manner. Furthermore, we have performed ChIP-seq in these cell lines to produce reproducible and high-resolution isoform-specific binding profiles for both SETα and SETβ. Integration of RNA-seq and ChIP-seq datasets reveals that expression of key players in the FGF signalling pathway are altered by SETα and SETβ addback. Interestingly, KLF5, a key factor regulating FGF signalling during primitive endoderm specification, uniquely binds the promoter of SETα and suppresses its expression. Together, these data hint at a relationship between SETα/SETβ alternative promoter switching at the KLF5-FGF signalling axis during early cell fate decisions.

## Results

### SETα is activated during major ZGA at the 2-cell stage

Initially, we investigated the expression of SET isoforms at the very first stages in development. We analysed published data from SMART-seq2 experiments in early mouse embryos which allow detection of alternative isoforms due to the use of full-length mRNA single-cell RNA sequencing (sc-RNAseq). Despite being expressed at low levels in ESCs, we found that SETβ transcripts cannot be detected at early and pre-blastocyst stages in vivo, while SETα transcripts can be detected from the 2-cell stage onwards (Fig. 1A).

**Figure 1.**
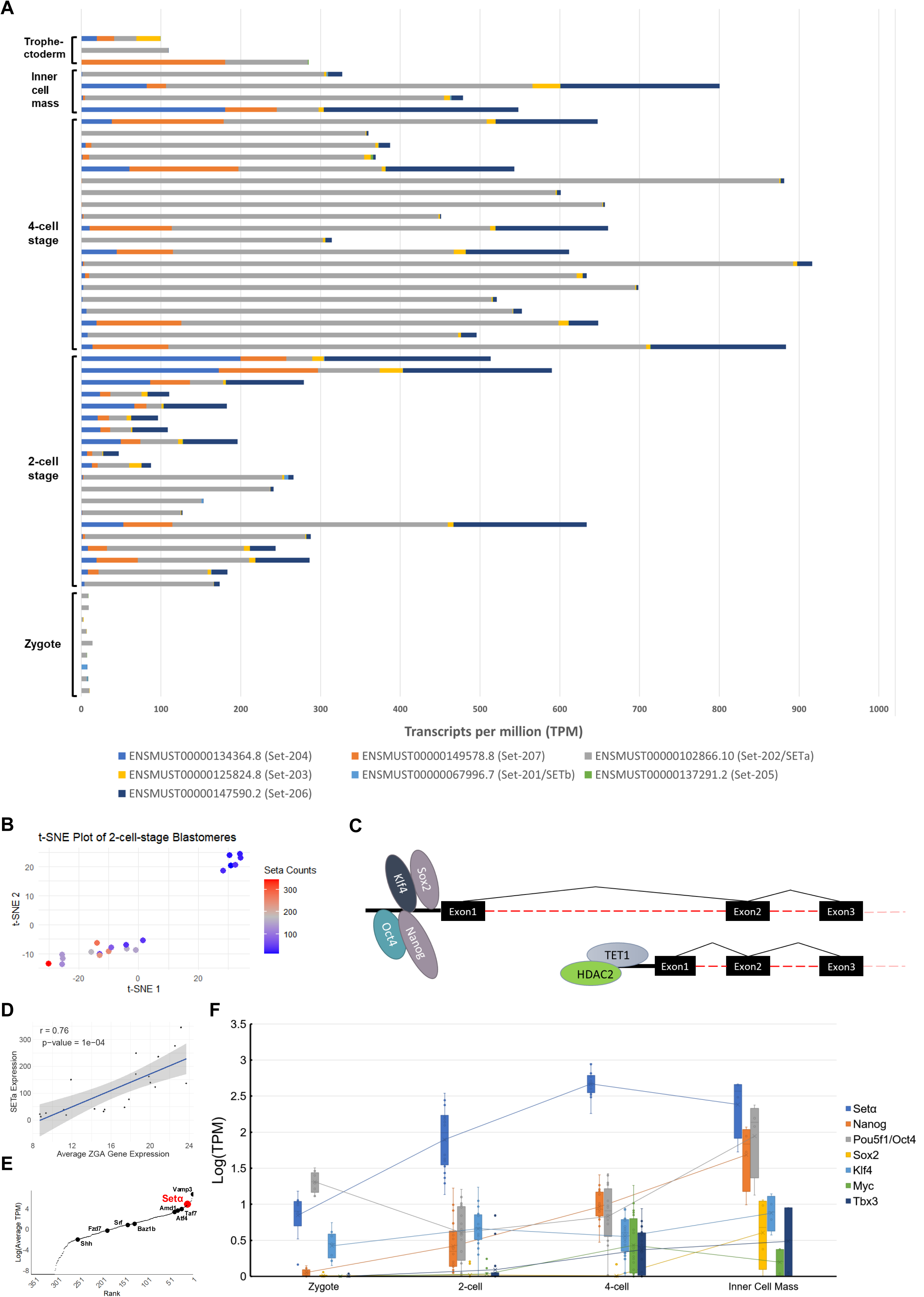
Expression of SET isoforms at the earliest developmental stages. A) Stacked bar chart comparing abundance (TPM) of different SET transcripts from sc-RNA-seq SMART-seq2 data in different pre-implantation stages of development. B) t-distributed stochastic neighbour embedding (t-SNE) plot and clustering of cells coloured by SETα transcript expression by transcript expression at the 2-cell stage. C) Binding of factors on SETα and SETβ promoters. D) Correlation between SETα transcript expression (TPM) and average major ZGA transcript expression (TPM). E) Ranked plot comparing average levels of SETα transcript expression in blastomere stage cells with transcripts activated during major ZGA. F) Levels of SETα expression (Log(TPM)) compared with pluripotency-associated factors at early stages of development.

Interestingly, although SETα is the predominantly expressed isoform at the 4-cell stage, 2-cell stage embryos appeared to be bifurcated into two sets of groups, SETα expressing cells and non-SETα expressing cells. In fact, clustering of cells at the 2-cell stage following dimensionality reduction and generation of t-SNE plots produces two distinct clusters of cells that appeared to segregate according to SETα expression (Fig. 1B). This suggests that SETα might itself be a marker, or otherwise co-expressed with a separate set of genes within a distinctive sub-two-cell stage identity. We previously revealed that differential expression of SETα and SETβ is regulated by distinct pools of TFs that may correspond to different nuclear contexts, with SETα being regulated by pluripotency factors (Fig. 1C) (Edupuganti et al., 2017; Svoboda, 2018). SETα may be activated prior to, or concurrent with, the establishment of the pluripotency.

Following fertilization and formation of the zygote, waves of transcription occur in a subset of zygotic genes, a phenomenon known as zygotic genome activation (ZGA). We therefore suspected that the bifurcation of SETα transcript expression levels was more likely to correspond to the first major ZGA, which is known to occur during the 2-cell stage (Hamatani et al., 2004). Looking at the average expression of genes activated during major ZGA, we observed a significant positive correlation (r = 0.76, Pearson’s correlation coefficient; P = 0.0001) between expression of the SETα isoform transcripts and ZGA genes (Fig. 1D), and non-significant weak correlations for other protein coding SET transcripts (Supp. Fig. 1A-C). This indicates that specifically SETα, rather than alternative SET isoforms, is among the transcripts activated in this manner in early-stage pre-implantation embryos. Moreover, of genes shown to be activated during major ZGA, SETα ranks among the top 5% of transcripts with the highest expression in blastomeres (Fig. 1E).

Interestingly, we observe that SETα’s activation at the 2-cell stage occurs earlier than other pluripotency-associated factors, including Nanog and Pou5f1 (also known as Oct4), which only become activated at similar levels in the inner cell mass (ICM) (Fig. 1F). Given SET’s role in maintaining a robust pluripotent state, this points to SETα being a zygotic activated gene that supports acquisition of pluripotency prior to activation of the core pluripotency factor network (Harikumar et al., 2020).

### SETα binding is enriched at genes linked to embryonic survival

To investigate isoform-specific chromatin binding, we performed ChIP-seq with HA antibodies after inducing either SETα-HA or SETβ-HA expression using our Dox-inducible system. Although we were able to retrieve transcription start site (TSS)-associated fragments following SET overexpression in WT ESCs, fewer than 130 of retrieved ChIP-seq peaks passed irreproducibility discovery rate thresholds across replicates (Supp. Fig. 2A). Using our Dox-inducible system, we immunoprecipitated only exogenously expressed SET fused with a HA-tag, which is in competition with endogenous SET for binding to chromatin. In addition, SET exists mainly as a dimer in the nucleus; SETα and SETβ homodimerize and can form heterodimers with each other; this limits the accuracy of detecting isoform specific binding in WT conditions (Miyaji-Yamaguchi et al., 1999; Muto et al., 2007). Thus, to detect isoform specific binding of SETα and SETβ, we sought ways to increase the quality of our SET-HA ChIP-seq experiments and uncover more reproducible peaks, as well as avoid the confounding factor of dimerization. Moreover, as mentioned above, commercially available antibodies that target the N-terminal region of SET are unavailable, which would have allowed for isoform-specific capture. To overcome these limitations, we performed ChIP-seq for SETα or SETβ following complete KO of endogenous SET in our KH2 SET Dox-inducible cell lines that express exogenous copies of either SETα or SETβ fused with HA-tag (Fig. 2A).

**Figure 2.**
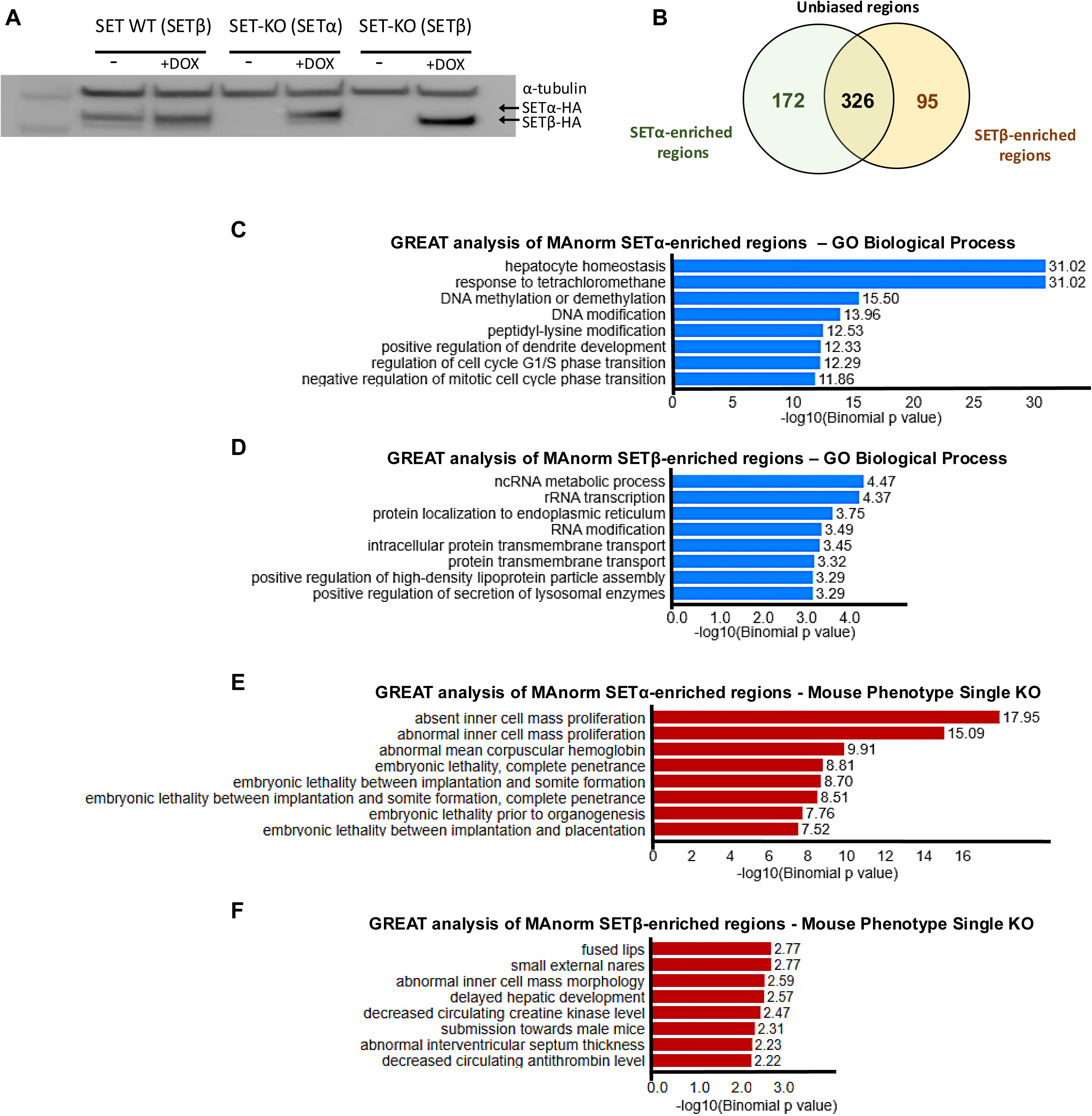
GREAT analysis of SETα and SETβ ChIP-seq regions. A) Western blot using anti-SET antibodies showing addback of SETα or SETβ in SET-KO background compared to SET-WT. α-tubulin (top band) was used as loading control. B) Venn diagrams comparing MAnorm-enriched IDR filtered MACS2-called peaks for SETα and SETβ. C) GREAT analysis of SETα-enriched regions for “GO Biological Processes” D) Same as (C) for SETβ. E) GREAT analysis of SETα-enriched regions for “Mouse Phenotype Single KO”. F) Same as (E) for SETβ.

Consequently, a pure population of SETα- or SETβ-bound DNA can be obtained from immunoprecipitation with HA antibodies following expression of SETα-HA or SETβ-HA. Reassuringly, we were able to observe a high reproducibility between MACS2-called peaks across two biological replicates. We obtained 530 and 485 peaks crossing the irreproducible discovery rate significance threshold (p < 0.05) for SETα and SETβ, respectively, of which 389 peaks overlapped between the two isoforms (Supp. Fig. 2B-C). In addition, most of SETα peaks (76.5%) were located within 5kb of TSSs (Supp. Fig. 2D-E) and we obtained considerably improved peak resolution (Supp. Fig. 2F).

Given the high degree of overlap between the two isoforms in their binding regions, we opted to perform a quantitative analysis to determine the relative strength of binding of each isoform compared with each other. To do this we used the MAnorm tool to normalize peaks between samples and compared the number of reads at each peak between our isoform-specific, SETα and SETβ ChIP-seq experiments (Supp. Fig. 3A). In doing so we obtained 172 regions specifically enriched for SETα and 95 regions specifically enriched for SETβ (Fig. 2B).

Using the Genomic Regions Enrichment of Annotations Tool (GREAT) (McLean et al., 2010), we functionally annotated genes associated with enriched SETα or SETβ regions we obtained with MAnorm. We generated terms unique to SETα or SETβ bound genes across three different databases, ‘GO Biological Process’, ‘GO Molecular Function’, and ‘Mouse Phenotype Single KO’. SETα preferentially targets genes relating to regulation of cell cycle progression as well as important genes for early embryo proliferation and survival, while both SETα and SETβ target genes were related to transcription, including genes involved in RNA polymerase II TF activity and sequence-specific DNA binding (Fig. 2C-F, Supp. Fig. 3B-C). This suggests that although both SET isoforms share redundancy in their target genes, SETα may possess a unique isoform-specific regulatory action on genes important for early embryonic processes.

### SETα and SETβ binding resembles ETS family of transcription factors

We used MEME-CHIP (Motif Analysis for Large Nucleotide Datasets) to analyse our SETα and SETβ IDR-filtered ChIP-seq peaks (Machanick and Bailey, 2011). With this analysis we detected the motif “CCGGAASY” as significantly enriched in SETα and SETβ overlapping target regions (Fig. 3A). The same analysis for total peaks called separately for both isoforms produced almost identical motifs. Indeed, for SETα and SETβ, 75% and 47% of peaks, respectively, matched these motifs, indicating high confidence that “CCGGAASY” is the consensus motif for SET (Supp. Fig. 3D-E). The closest matches to this motif are the binding motifs of ETV4 and ELK4, members of the highly evolutionarily conserved ETS family of TFs (Fig. 3B).

**Figure 3.**
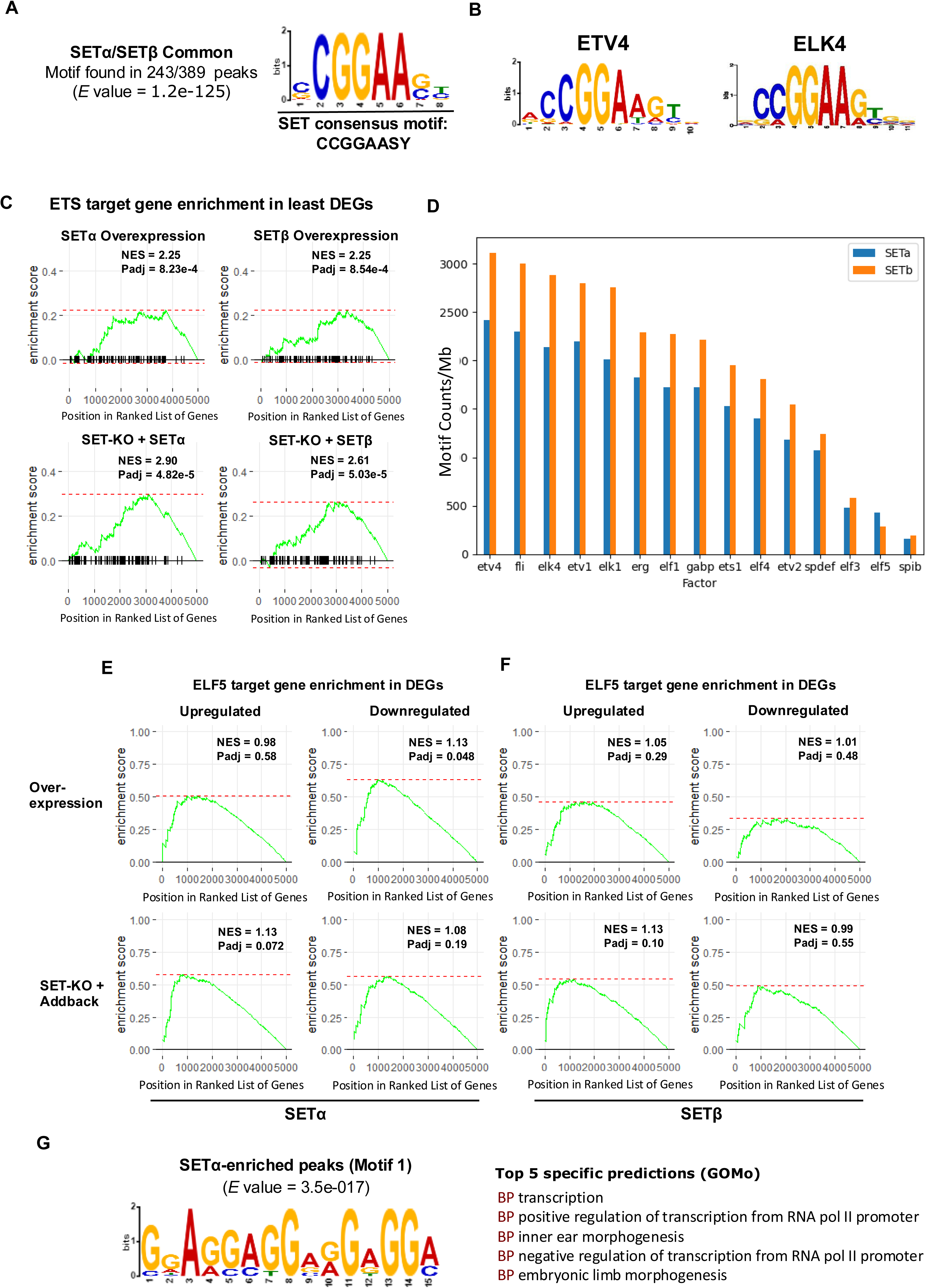
Predicted Motifs for SETα and SETβ are related to ETS family factors. A) Predicted binding motif for SETα/SETβ overlapping peaks as identified by de novo motif discovery analysis using the motif-based sequence analysis tool MEME-ChIP. B) Binding motif for ETS domain -containing proteins ETV4 (top) and ELK4 (bottom). C) GSEA plots displaying enrichment of ETS family target genes based on least differentially expressed genes from SETα/SETβ overexpression (top) and SET-KO with SETα/SETβ addback (bottom). D) Abundance (counts per megabase) of ETS family motifs in peaks from SETα and SETβ ChIP-seq data. E) GSEA plots displaying enrichment of ELF5 target genes based on upregulated (left) or downregulated (right) genes from SETα overexpression (top) and SET-KO with SETα addback (bottom). F) Same as (E) for SETβ. G) Top significant MEME-ChIP predicted de novo motif for MAnorm-enriched SETα regions (left) and associated GOMo predicted regulatory gene ontology terms (right).

Members of the ETS family are implicated as being highly active in tumorigenesis due to pro-oncogenic ETS signalling. ETS factors are defined by the possession of an ETS-domain that enables DNA-binding and they bind a core purine-rich 5lll-GGAA/T-3lll motif (Sharrocks, 2001; Sizemore et al., 2017). As SET does not contain an ETS-domain, it is possible that SET participates in a complex with ETV4/ELK4 or other ETS factors. However, from our mass spectrometry data, we did not detect an interaction of either SETα or SETβ with ETS family proteins. SET may therefore share redundant roles with ETS factors in the regulation of ETS target genes, many of which are essential for embryonic development. ETS factors are known for having versatile functions and can act as activators or repressors under different cellular contexts (Kar and Gutierrez-Hartmann, 2013). Importantly, 11 out of the 27 murine *Ets* genes have been shown to result in complete or significant embryonic and/or postnatal lethality in experiments with knockout or mutant mice (Findlay et al., 2013).

We next wished to see if SET could regulate ETS target genes. To accomplish this, we performed overexpression of SETα or SETβ and looked at differential gene expression following RNA-seq. In addition, to account for the existing presence of SETα/ SETβ in WT ESCs, we repeated the experiment, this time performing RNA-seq following SETα or SETβ addback in a SET-KO background. Using publicly available data (Hollenhorst et al., 2011), we obtained a list of ETS target genes by considering genes that were bound at their promoters by at least two of the ETS factors, as detected by ChIP experiments. This gave us a list of 863 genes which we mapped to 250 mouse genes with SynGO (Koopmans et al., 2019). With gene set enrichment analysis (GSEA) analysis we looked at the enrichment of ETS target genes based on their expression following SET overexpression or SET addback following SET KO. We did not observe a strong enrichment of ETS target genes in differentially expressed genes (DEGs) following individual SET isoform overexpression or addback. Only a handful of ETS targets were among the top 5000 DEGs for both up- and down-regulated gene lists ranked by log2 fold change. GSEA enrichment scores (ES) were < 0 for all groups of DEGs, indicating underrepresentation of ETS targets (Supp. Fig. 4A-B). However, interestingly, looking at the DEGs least differentially expressed revealed high significant enrichment (Fig. 3C). This suggests that although SET appears unable to directly regulate ETS factors target genes, SET may be preferentially regulating genes that are not bound by ETS factors.

To determine if there were specific factors in the ETS family that might share more similarity to either SETα or SETβ, we calculated the abundance of 15 ETS factors with established and available motifs (Heinz et al., 2010). The most abundant motif for both SETα and SETβ, was that of ETV4. Like SETα, ETV4 is regulated by OCT3/4 and supports proliferation in ESCs (Akagi et al., 2015). All ETS factor motifs were more abundant in SETβ peaks compared with SETα with the exception of ELF5 (Fig. 3D). Interestingly, ELF5 has been implicated in early blastocyst cell fate, during specification of the trophectoderm, a stage where SETβ is not expressed (Fig. 1A). Performing GSEA on ELF5 target genes obtained from JASPAR Predicted Transcription Factor Targets (Rouillard et al., 2016) revealed a significant enrichment (Padj = 0.048) of ELF5 targets in downregulated genes following SETα overexpression in WT conditions (but not following SETβ overexpression), which was not the case following GSEA on ELK4 target genes (Supp. Fig. 4C-D). These results suggest that SETα may possess an isoform-specific and selective repressive function on ELF5 targets.

We next performed MEME-ChIP analysis on our list of SETα/SETβ MAnorm-enriched regions. Interestingly, we obtained a unique significantly enriched motif for SETα (Fig. 3G, left). For both SETα- and SETβ-enriched regions we obtained similar significant motifs resembling ETS family factors, however, we detected that SETα is more sensitive to T/C at the 8^th^ bp position of the SET consensus “CCGGAASY” motif (Supp. Fig. 5A-B). After performing gene ontology for motifs (GOMo) analysis for significant motifs for SETα/SETβ we revealed that the SETα-specific motif has additional regulatory functions that include several developmental-related processes (Fig. 3G, right). GSEA for genes with promoters that contain this motif do not display significant enrichment although the enrichment trend between conditions resembles our ELF5 target gene GSEA results (Fig. 3E-F, Supp. Fig. 5C-D). Overall, these data show that both SET isoforms strongly resemble the binding profile of ETS family proteins but seem to preferentially regulate genes that are selectively unbound by ETS factors. Additionally, SETα may have an additional ability to bind and regulate a wider array of target genes, and in particular targets of ELF5.

### SETα and SETβ differentially regulate their target genes

To determine if any of the isoform-specific target genes we identified in our ChIP-seq assays could be regulated by SET in WT conditions, we looked at the expression of these genes in our SETα/SETβ overexpression and addback RNA-seq data. In WT ESCs we found that SETβ negatively regulates its target genes, while SETα has minimal effect. However, as SETα is predominantly expressed in WT ESCs, overexpression of SETα might have minimal effect on its target genes if they are already expressed at saturation levels. KO of both SET isoforms leads to downregulation of both SETα and SETβ target genes. Additionally, comparing DEGs from SETα-KO and SETβ-KO datasets, we found that SET depletion in both cases leads to upregulation of common SET targets with minor effects on SET-isoform specific targets (Fig. 4A). This indicates that both SET isoforms have an overall negative regulatory effect on their shared targets.

**Figure 4.**
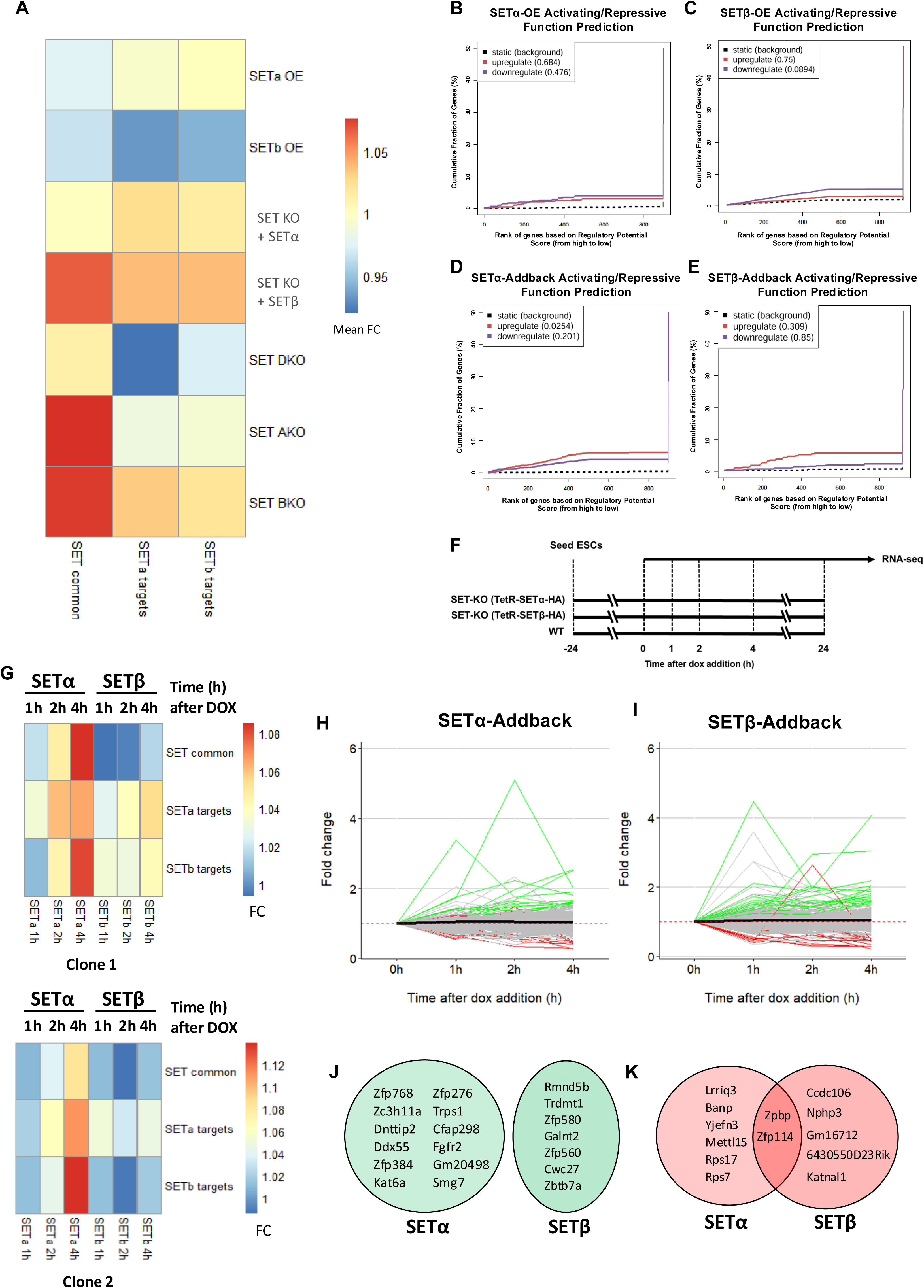
Regulatory function of SETα and SETβ at target genes. A) Heatmap of RNA-seq data showing mean expression fold change for SET common and isoform-specific targets in different SET expression conditions. B) BETA functional prediction for SETα targets and differentially expressed genes following SETα overexpression. Genes are ranked according to predicted regulatory potential score on the X-axis. The Y-axis indicates fraction of genes in the upregulated/downregulated gene group that have a higher regulatory potential score for their respective rank. Upregulated and downregulated genes are indicated by red and purple lines, respectively. Dotted lines represent non-differentially expressed genes which are used as background. P values represent the significance of the upregulation/downregulation group compared to non-differentially expressed group using the Kolmogorov-Smirnov test. C) Same as (B) for BETA functional prediction for SETβ targets and differentially expressed genes following SETβ overexpression. D) Same as (B) for BETA functional prediction for SETα targets and differentially expressed genes following SETα addback in SET-KO cells. E) Same as (B) for BETA functional prediction for SETβ targets and differentially expressed genes following SETα addback in SET-KO cells. F) Schema for SETα and SETβ addback time-course RNA-seq experiments. G) Heatmap of foldchanges of mean expression for SET common and isoform-specific targets after 1h, 2h and 4h of Dox treatment in two independently-derived clones. H) Trajectory of expression of genes that have upwards or downwards trajectory after 4h of Dox treatment for SETα-addback. Foldchange is compared to the 0 h timepoint after normalization by WT expression. I) Same as (H) for SETβ. J) List of SETα and SETβ ChIP-seq target genes that show upwards expression trajectory after 4h Dox treatment. K) Same as (J) for target genes that show downwards expression trajectory.

To identify more general patterns in SET’s regulatory function on its target genes, we performed binding and expression target analysis (BETA) and looked at predicted targets based on peaks within 3kb of TSSs of DEGs. BETA predicts target genes of factors and regulatory potential by combining ChIP-seq binding data with differential expression data (Wang et al., 2013). Overexpression of SETα in WT ESCs did not have substantial effects on gene expression, however SETβ overexpression had a significant predicted repressive function on its targets (p = 0.089) (Fig. 4B-C). Interestingly, in a SET-KO background, we found that SETα addback was able to significantly activate (p = 0.025) and non-significantly downregulate (p = 0.201) its target genes, and a non-significant activating (p = 0.309) and repressive (p = 0.85) function for SETβ addback (Fig. 4D-E).

To investigate more specific and immediate effects on gene regulation, we next performed time-course addback RNA-seq experiments over a period of 4 hours (Fig. 4F). We used K-means clustering of our time-course RNA-seq data to obtain 5 clusters of genes that showed similar patterns of expression. We repeated this experiment with another independently derived clone and generated a pairwise heatmap to compare cluster similarity and obtained high-confidence clusters between two clones that shared more than 50% of genes (Supp. Fig. 6A-B). We then took the top four high confidence sets of genes for SETα and SETβ and performed KEGG enrichment analysis, which revealed known functions of SET including cell-cycle regulation, p53-signalling, and cancer related functional terms (Supp. Fig. 7).

In confirmation of our BETA regulatory function prediction (Fig. 4B-E), we find that SETα can activate SET targets more effectively than SETβ, with a similar trend observed in two independently derived clones (Fig. 4G-H). Next, we looked at individual genes that showed an upwards or downwards trajectory in gene expression over 4 hours of SETα or SETβ addback. We looked at whether these genes were also targets of SET from our ChIP-seq experiments and identified groups of SETα/SETβ-activating and SETα/SETβ-repressing genes (Fig. 4J-K). Among the SETα activating genes were *Zc3h11a*, the protein product of which was shown to bind mRNA of metabolic genes important for embryonic survival (Younis et al., 2023), *Trps1*, which is important for mouse preimplantation embryo development, as well as fibroblast growth factor receptor, *Fgfr2*, which, together with *Fgfr1*, is essential for primitive endoderm (PrE) formation (Molotkov et al., 2017) (Fig. 4J). As each of these genes is bound by both SETα and SETβ, this suggests that SETα-specific protein complexes may be involved regulating unique SETα-activating genes (Supp. Fig. 8A-C).

### SETα repression by KLF5 occurs during blastocyst development

Interestingly, in a previous study we found that SET KO in ESCs led to upregulation of PrE markers, which could not be rescued following differentiation (Harikumar et al., 2020). Since we detected *Fgfr2* as potentially regulated by SET, we compared other central factors involved in FGF signalling during PrE specification. In our time-course add-back experiments, we find that SETα upregulates *Fgfr1*, *Fgfr2, Fgf4* and *Nanog* (Fig. 5A), while *Fgf4* is downregulated by SETβ (Fig. 5B). During PrE specification, FGF4 levels are important for specifying blastocyst cell identities. Namely, FGF4 activation of FGFR1/FGFR2 determine the extent of Erk signalling which primes blastomeres to become either the pluripotent epiblast or PrE in the late blastocyst (Hermitte and Chazaud, 2014). Interestingly, we observe binding of both SET isoforms at the *Fgf4* locus, with a slight enrichment of SETα, although this was not identified by our MAnorm analysis (Supp. Fig. 8D). It is possible that differential regulation of *Fgf4* by SETα and SETβ may be the result of isoform-specific binding of different sets of co-regulatory factors.

**Figure 5.**
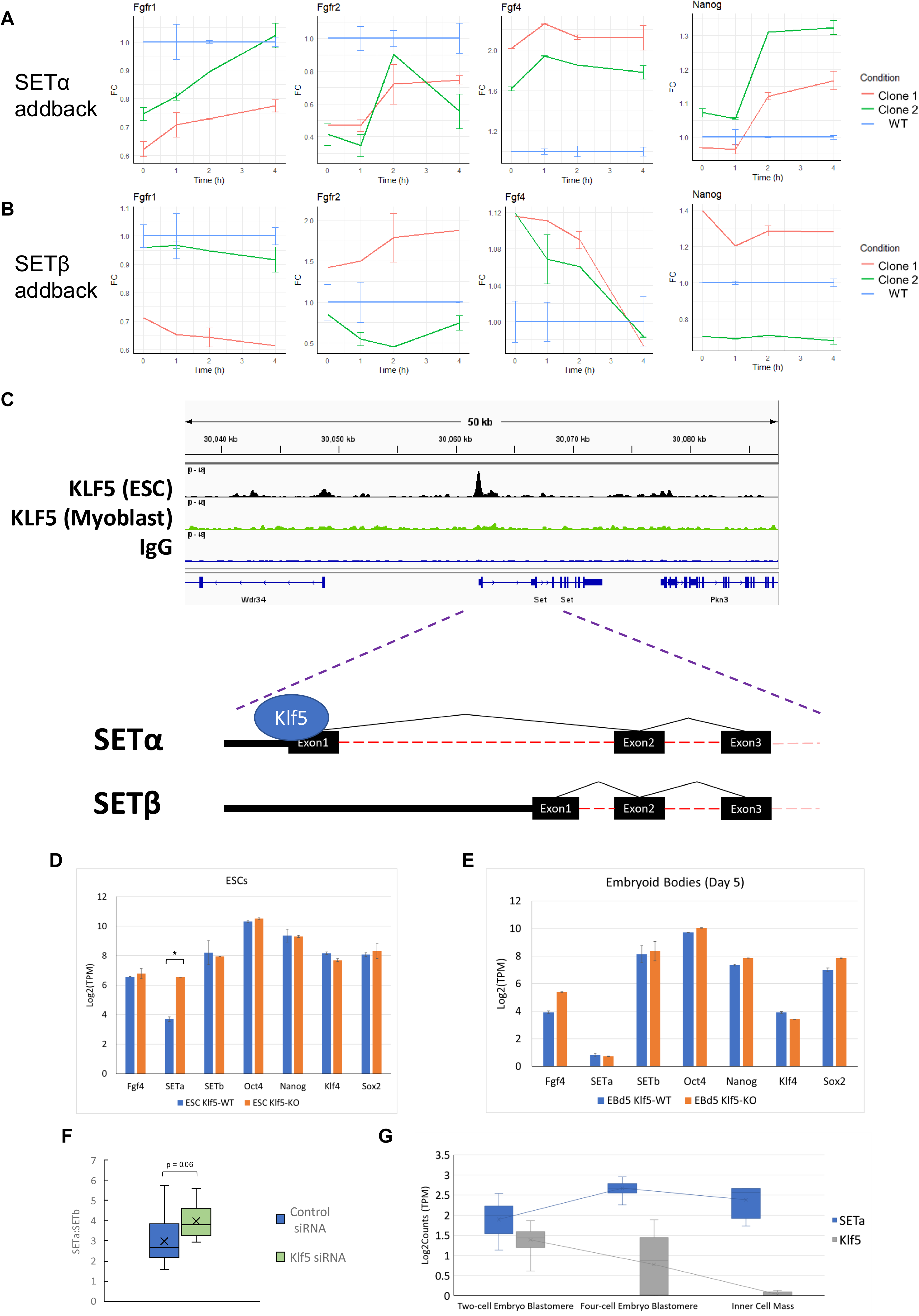
Regulation of FGF-signalling genes by SET. A) Expression changes in two clones for genes important for FGF signalling during primitive endoderm specification following SETα addback in SET-KO ESCs. B) Same as (A) for SETβ addback. C) IGV bigwig track for KLF5 ChIP-seq data in ESCs (red) and myoblasts (green) at the SETα/SETβ gene locus. D) Expression of SETα, SETβ, and other pluripotency factors following *Klf5* KO in ESCs E) Same as (D) in embryoid bodies. F) Ratio of SETα to SETβ expression levels following *Klf5* knockdown with siRNA compared to WT. G) Levels (Log2 TPM) of *Setα* compared with *Klf5* at the two-cell stage, four-cell stage, and early blastocyst inner mass embryonic stages.

Another important factor involved in PrE specification is KLF5. *Klf5* KO embryos have elevated levels of FGF4, leading to adoption of a PrE lineage fate. Correspondingly, KLF5 overexpression represses FGF4, resulting in NANOG persistence and incomplete segregation of PrE fate (Azami et al., 2017). As we detected lower levels of *Fgf4* following SETβ addback, we wondered if KLF5 might act upstream of SET. Therefore, we next analysed publicly available KLF5 ChIP-seq data and RNA-seq data from KLF5-depletion experiments. Surprisingly, we found that in ESCs, KLF5 specifically binds the promoter of *Setα*, but not of *Setβ* (Kinisu et al., 2021) (Fig. 5C, red tracks). This binding is specific to ESCs as, interestingly, we do not observe KLF5 binding of the *Setα* promoter in mouse C2C12 myoblasts (Hayashi et al., 2016) (Fig. 5C, green tracks). Using publicly available data from the Gene Expression Omnibus (GEO Accession Number: GSE202540) we found that *Klf5* KO experiments in ESCs show a selective induction in SETα expression (Fig. 5D). This was not the case following differentiation into embryoid bodies (Fig. 5E), where, as expected, SETα expression is significantly diminished, again demonstrating an ESC-specific relationship between KLF5 and SET. In siRNA experiments the same trend can be observed, with a greater ratio of SETα to SETβ levels upon *Klf5* knockdown (Fig. 5F) (Kinisu et al., 2021). This indicates that KLF5 has an ESC-specific regulatory effect on the expression of SETα, but not SETβ.

Finally, comparing levels of KLF5 and SETα in blastomeres and in the blastocyst inner cell mass, we observe a decrease in KLF5 expression that corresponds to an increase in SETα expression (Fig. 5G). Taken together, these data support a model by which KLF5 represses SETα but not SETβ expression in a manner temporally restricted to early embryonic development.

### SETα has greater predicted intrinsic disorder in its N-terminus

How might isoform-specific regulation be encoded by differential SETα/SETβ protein sequence? Structurally, SETα and SETβ differ in short N-terminal regions which are coded for by alternative first exons. This N-terminal region is 36 and 24 amino acids in length for SETα and SETβ isoforms, respectively. We used the deep learning protein structure prediction method, Alphafold2, to approximate the protein structure of both SET isoforms. SETα appears to display a high degree of intrinsic disorder in its N-terminus (Fig. 6A, green shade), while SETβ has a predicted ordered alpha helix conformation (Fig. 6B, yellow shade).

**Figure 6.**
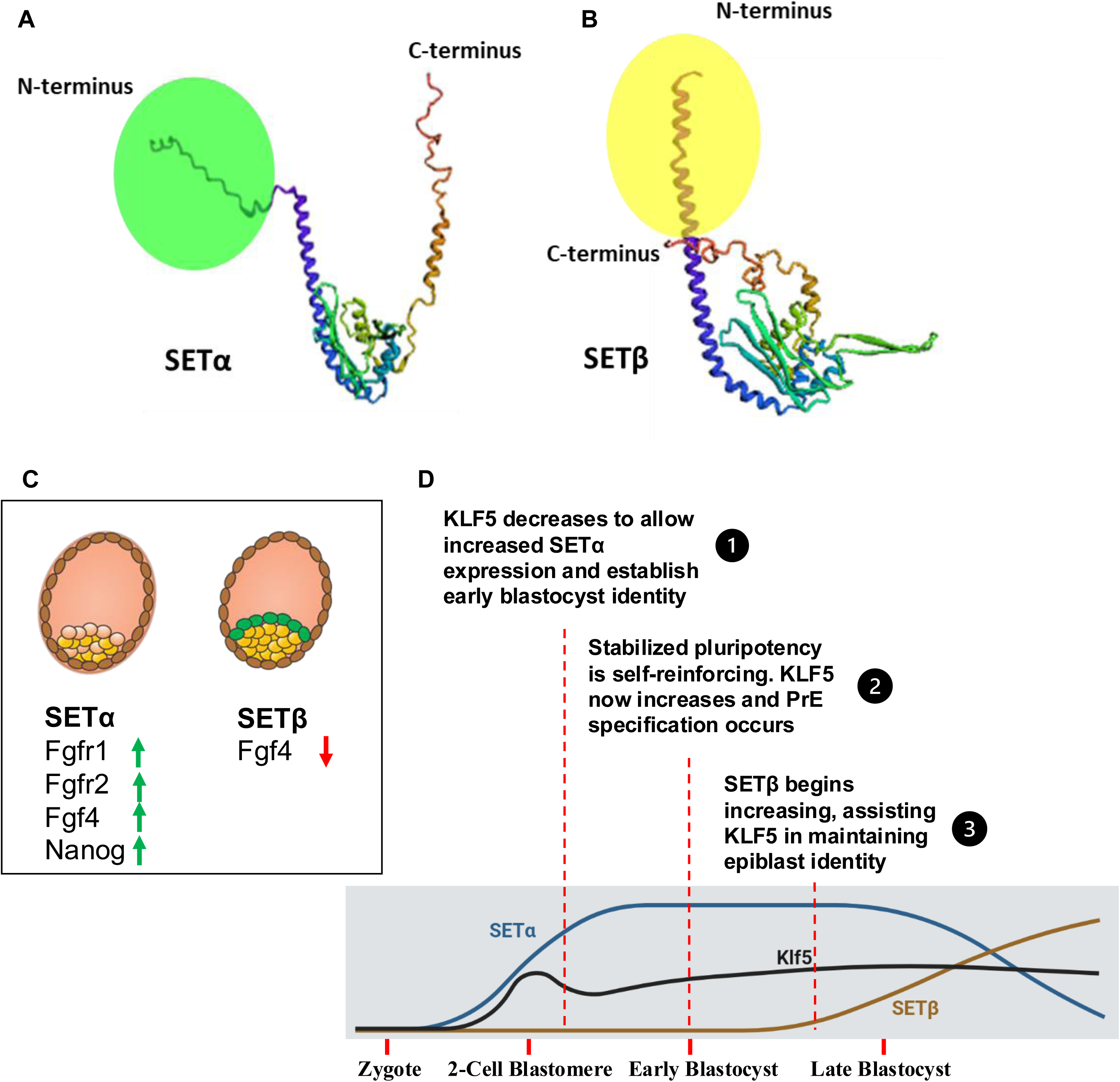
Hypothesis of SET isoform function in the pre-implantation embryo. A) Alphafold2 predicted structure for SETα. Green circle highlights the SETα-specific N-terminus. B) Alphafold2 predicted structure for SETβ. Yellow circle highlights the SETβ-specific N-terminus. C) Expression changes of important players in FGF-signalling in the early vs late blastocyst. D) Model describing how expression dynamics of SETα/SETβ/KLF5 in the early embryo may play a role in primitive endoderm specification and epiblast identity.

The increased number of SETα-specific ChIP-seq peaks (Fig. 2B, Supp. Fig. 2C) may be reflective of the disordered N-terminus of SETα, which may extend out from SET’s dimerization domain and facilitate a greater diversity of chromatin-binding interactions (Muto et al., 2007). Moreover, the lower abundance of ETS motifs in SETα peaks may be explained by a more promiscuous binding to chromatin due to the greater intrinsic disorder of SETα. Indeed, intrinsic disordered regions (IDRs) in non-DNA binding domains in TFs have been implicated in wider specificities of promoter selection (Kumar et al., 2023). This is supported by a reduced overall abundance of ETS motif counts at SETα peaks (Fig. 3D), which may potentially serve to dilute SETα at ETS target sites and encourage SETα’s participation in protein complexes with different chromatin specificities.

To determine how this difference in disorder might influence protein binding, we reanalysed mass spectrometry data and included SET isoform specific KO experiments. We obtained lists of proteins that could be recovered in two replicates for SET-WT as well as for SETα-KO and SETβ-KO only samples. Entering these lists of proteins into the STRING database we found several complexes or protein families that were specific to SET-WT, which would include both SET as a heterodimer or as a homodimer (Szklarczyk et al., 2023). In addition, using data from SETα-KO and SETβ-KO samples, we detect proteins that bind only one of the two SET-isoforms (Supp. Fig. 9A). In theory, intrinsic disorder engenders higher conformational flexibility, allowing a greater degree of promiscuous binding (Babu, 2016), although we observed that for SETβ (in SETα KO ESCs) we obtained a higher abundance of interacting proteins than for SETα (in SETβ KO ESCs) (Supp. Fig. 9B). However, this may be due to the absolute levels of SET protein, as knockout of SETα leads to increased expression of SETβ in ESCs (Edupuganti et al., 2017). Unique to SETα were several proteins involved in chromatin remodelling (members of Ino80 and SWI/SNF complexes) (Supp. Fig. 9C), unique to SETβ were H1 histone proteins (HIST1H1a, HIST1H1C, and HIST1H1E) (Supp. Fig. 9D), and proteins bound only to SET heterodimers in WT cells included ribosomal-associated proteins (RCL1 and RRP9) (data not shown). It is feasible that an association with chromatin remodelling factors may confer the SETα’s unique regulatory functions revealed in this study.

Overall, our study identifies isoform-specific regulatory functions for SETα/SETβ. The appropriate balance of SET isoform expression levels may be important to ensure correct and timely cell-fate decisions in the early blastocyst.

## Methods

### Cell culture

KH2 mouse ESCs were cultured using standard procedures on mouse embryonic fibroblast (MEF) coated plates, in DMEM (Sigma) supplemented with 15% fetal bovine serum (FBS) (Biological Industries), L-glutamine, 1X MEM non-essential amino acids, Sodium Pyruvate, 1x Penstrep, 0.2 mM β-ME and leukemia inhibitory factor (LIF). Cell cultures were maintained in a humidified atmosphere (5% CO2 at 37°C). Depletion of MEFs was performed by pre-plating trypsinized ESCs on low attachment plates twice, for 30 mins each. KH2 mouse ESCs were used to create SET-OE cell lines with SETα-HA or SETβ-HA expression under the control of a tetR Dox-inducible promoter using the pBS31 Vector (Open Biosystems). Although e-karyotyping did not detect any abnormalities, cytogenetic karyotyping of our KH2 clones revealed potential duplications in chromosome 8 and 11. Therefore, to ensure duplication-based bias, chromosome 8 and 11 was excluded from analyses involving KH2 ESCs. Overexpression constructs were introduced by co-transfection with Flp recombinase using LT1 transfection reagent (Mirus). Selected KH2-SETα and KH2-SETβ clones were expanded and maintained on low hygromycin (100 µg/mL). Overexpression assays were carried out with 1 μg/mL of Dox in mouse ESC media. CRISPR KO in KH2 ESCs were generated with guide RNAs were targeting exon 4 of SET and exons 1 of SETα or SETβ in R1 ESCs, as previously described (Edupuganti et al., 2017; Harikumar et al., 2020; Ran et al., 2013). CRISPR/Cas9 target sequences used were as follows: TCAGCCCAAGAAACCCAGACCGG (for generating KO of SETα), GGGCACCATGTCTGCGCCGACGG (for generating KO of SETβ), CCCACCACATACCTTGTGGATGG (for generating KO of both SET isoforms).

### Western blots and antibodies

Confluent ESCs were harvested following MEF depletion by pre-plating trypsinized cells on low attachment plates twice, for 25 mins each. Cells were washed with PBS, trypsinized and collected by mild centrifugation. Cells were then resuspended in RIPA buffer containing protease inhibitor cocktail. Lysates were incubated on ice for 30 min and centrifuged to remove cell debris. The resulting supernatant was subjected to further analysis. The following antibodies were used in this study: SET (A302-261A, Bethyl), α-TUBULIN (ab4074, Abcam) and HA (ab9110, Abcam).

### Analysis of SMART-seq2 datasets

Raw sequencing data from SMART-seq2 experiments was obtained from NCBI (GSE57249), which consisted of single-cell RNA-seq data for 9 zygotes, 10 2-cell, and 5 4-cell mouse (C57BL/6) embryos (Biase et al., 2014). Reads were aligned and quantified according to transcript-level abundance with Salmon (v1.9.0) (Patro et al., 2017). Dimensionality reduction and cluster generation was performed using the t-SNE method in R.

### SET isoform structure comparison

Deep learning protein prediction method, Alphafold2, was used for predictions of SETα and SETβ protein structure (Jumper et al., 2021).

### RNA extraction and sequencing

RNA extraction was performed using RNeasy Mini Kit (Qiagen). 500ng of RNA was used to generate libraries with QuantSeq 3’ mRNASeq Kit (Lexogen) according to the manufacturer’s instructions. RNA-seq libraries were sequenced in-house on NextSeq™ 2000, or Illumina HiSeq 2500 machines. Sequencing reads from RNA-seq experiments were aligned with STAR (v2.5.4b) (Dobin et al., 2013) using default settings against the mm10 mouse genome obtained from ENSEMBL and reads were counted with featureCounts (Liao et al., 2014). Differential expression analysis was performed with DESeq2 (Love et al., 2014) and the DESeq2 counts function was used to obtain sizefactors for each sample to calculate normalized count comparisons. Gene expression heatmaps, boxplots and density plots of log2 fold expression changes were produced using the R packages, ggplot2 and plyr. Expression data from KLF5 KO (Accession Number: GSE202540) and siRNA (Accession Number: GSE186005) experiments (Kinisu et al., 2021) were obtained from publicly available GEO datasets and were processed in the same manner as the RNA-seq data generated in this study.

### Chromatin immunoprecipitation followed by sequencing (ChIP-seq)

Following MEF depletion, 10 million ESCs were harvested and crosslinked with 1% formaldehyde for 10 min and reaction was quenched with glycine. DSG was added at a final concentration of 2 mM for 45 min at RT prior to addition of formaldehyde. Cells were lysed at 4°C for 10 minutes in lysis buffer 1 (50 mM HEPES-KOH 1 mM EDTA, 140 mM NaCl, 0.25% Triton X-100, 0.5% NP-40, 10% glycerol) 10 min in lysis buffer 2 (1 m mM Tris-HCl, 1 mM EDTA, 200 mM NaCl and 0.5 mM EGTA), and 30 min in lysis buffer 3 (10 mM Tris-HCl, 1 mM EDTA, 100 mM NaCl, 0.5mM EGTA and 0.1% Na-DOC). All lysis buffers were supplemented with protease inhibitors. Sonication was performed with Bioruptor® (Diagenode) for 27 cycles (30s on, 30s off), or until average DNA fragment size ranged from 300-500bp. Average sizes of libraries were determined by TapeStation (Agilent Technologies, USA). Samples were then incubated overnight at 4°C with Magna ChIP™ protein A beads (Sigma-Aldrich) coupled with 3 μg of HA (ab9110, Abcam). The beads were washed twice at 4°C with ice-cold RIPA buffer (10 mM Tris-HCl, 1mM EDTA, 140 mM NaCl, 1% Triton X-100, 0.1% SDS, and 0.1% Na-DOC) and once with RIPA-500 buffer (10 mM Tris-HCl, 1 mM EDTA, 500 mM NaCl, 1% Triton X-100, 0.1% SDS, and 0.1% Na-DOC), LiCl wash buffer (10mM Tris-HCl, 1 mM EDTA, 250 mM LiCl, 0.5% NP-40, and 0.5% Na-DOC), TE buffer (10 mM Tris-HCl and 1 mM EDTA), and eluted from the beads in direct elution buffer (10 mM Tris-HCl, 5 mM EDTA, 300 mM NaCl, and 0.5% SDS). The supernatant was reverse-crosslinked with RNase and 1U Proteinase K per sample for at least 4 h at 65°C. Reverse-crosslinked DNA was purified with AMPure XP SPRI beads (Beckman). Quantification of DNA concentration was performed with Qubit fluorometric quantitation (Thermo Fisher Scientific). Library preparation was carried out as previously published (Blecher-Gonen et al., 2013). Reads were sequenced in-house on an Illumina NextSeq™ 2000 machine. ChIP-seq analysis was performed following alignment of reads with bowtie2 (Langmead and Salzberg, 2012) to the GRCm38 (mm10) genome and peaks were called with MACS2 (Feng et al., 2012). Bigwig files were generated from aligned and sorted BAM files using bamCoverage (deeptools) (with -e 250 -- normalizeUsing RPGC). MAnorm (Shao et al., 2012) (--s1 100 --s2 100 -w 1000 -m 0.2 -p 0.05) was used to quantitatively compare peaks between ChIP-seq experiments. Reads within ENCODE blacklisted regions were filtered out prior to analyses. Supporting KLF5 ChIP-seq data was obtained from obtained from the NCBI SRA database (GEO accession: GSE137036 (for ESCs) and GSE80812 (for myoblasts)) and was processed in the same manner as the ChIP-seq data generated in this study (Hayashi et al., 2016; Kinisu et al., 2021).

## Discussion

Separate pools of TFs are targeted to two different promoters that regulate the expression of SETα and SETβ during early embryonic development. Although alternative promoter usage is thought to increase transcriptome and proteome diversity, it has been argued that molecular error is responsible for the majority of multiple alternative transcription initiation events at protein-coding genes, suggesting that most alternative isoform expression is nonadaptive (Xu et al., 2019). However, as the conversion from SETα to SETβ expression occurs at a defined developmental transition (i.e. pluripotency exit), this hints at a conserved regulatory function of SET isoform switching. Nevertheless, it remains difficult to establish the extent which this transcriptional switch is relevant for proper development of the embryo.

Our analysis of SMART-seq2 scRNA-seq data reveals that SETα is constitutively expressed from the 2-cell stage onwards, well into formation of the early blastocyst. SETβ, is however, absent at these stages, despite being expressed in cultured ESCs—albeit at a lower level than SETα. The lack of SETβ at these early stages suggests either that SETβ is only important for a later developmental program and may interfere with earlier developmental stages, or that SETα is important for supporting pre-blastocyst regulatory programs. However, as SET-KO mouse embryos survive until embryonic day 12.5, SET may be dispensable at these initial post-zygotic stages (Kon et al., 2019).

The relatively higher expression of SETα compared with other SET isoforms in early pre-implantation stages may be explained by the higher levels of pluripotency factors, which are expressed from maternally provided transcripts or otherwise activated at the 2-cell stage during major ZGA (Goolam et al., 2016; Svoboda, 2018). This agrees with our previous experiments where we demonstrated that the SETα promoter is bound by pluripotency factors (Edupuganti et al., 2017). However, levels of SETα are higher than most pluripotency-associated factors at pre-blastocyst stages, suggesting that SETα may be supporting cellular programs upstream of pluripotency acquisition.

Expression levels of SETα correlate with the expression of genes activated during the first major wave of ZGA, which occurs to activate the zygotic genome following decay of maternally provided RNA and peptides. Indeed, we found that SETα is among the highest expressed ZGA transcripts in mouse blastomeres. Transcripts activated in this fashion are involved in metabolic and housekeeping function and include ribosomal and RNA-processing related genes (Svoboda, 2018). Interestingly, we found that many of these genes are also regulated by SET, suggesting that activation of SETα may be important for embryonic survival prior to pluripotency establishment.

To identify potential regulatory processes the two SET isoforms might be uniquely involved in, we focused on the transcriptional response to SET expression, its binding profile on chromatin, and investigated potential isoform-specific regulation of expression by SETα or SETβ. From ChIP-seq experiments we obtained almost identical motifs for both SETα and SETβ which resembled ETS family members, ETV4 and ELK4. As ETV4/ELK4 or other ETS family members were not present in SET immunoprecipitation followed by mass spectrometry experiments, SET may not be cooperatively binding with ETS factors, suggesting that SET may interfere with ETS factor binding, or function on a similar set of genes and share a similarity in motif specificity. Interestingly, however, we found that SETα ChIP-seq peaks contained a higher abundance of ELF5 binding motifs and that SETα but not SETβ possesses a repressive function on ELF5 target genes. As this occurs only in WT conditions, this points to a pluripotency-stage specific regulatory role for SETα. In SET-KO conditions, ESCs begin to exit the pluripotency state which may preclude detection of SETα’s regulatory function in early embryonic stages (Harikumar et al., 2020).

Integration of our ChIP-seq data, specifically SET-isoform specific bound genes and RNA-seq data following SETα/SETβ overexpression and SETα/SETβ addback, revealed subtle activation or repressing functions as well as direct regulatory effects on target genes. We identified SET target genes whose expression is specifically regulated by either SETα or SETβ. Several of these targets are implicated in the FGF signalling pathway which is important during primitive endoderm specification.

In the late morula, heterogeneity in FGF4 levels results in a “salt-and-pepper” distribution which leads to segregation of the primitive endoderm from the inner cell mass of the early blastocyst (Azami et al., 2017). Interestingly, in previous work we demonstrated that although markers of all three germ layers are upregulated following *Set* KO, markers of primitive endoderm are not rescued after 4 days of retinoic acid differentiation (Harikumar et al., 2020).

Based on our current investigation, we have delineated potential isoform-specific roles for SETα and SETβ. Specifically, that the dynamic transcriptional activation of initially SETα, occurring prior to establishment of pluripotency, and subsequently, activation of SETβ following stabilized pluripotency in the blastocyst, contributes to maintaining epiblast and primitive endoderm identity. SETα promotes expression of factors that promote primitive endoderm/epiblast segregation such as FGFR1, FGFR2, FGF4 and NANOG, while SETβ negatively regulates FGF4. An important regulator of FGF4 expression is KLF5, whose absence skews the cellular proportions of primitive endoderm and epiblast in the late blastocyst (Azami et al., 2017). As KLF5 binds the promoter of SETα specifically in ESCs and represses its expression, we therefore suggest that KLF5—which is activated by Dux and other 2C-specific TFs in the 2-cell blastomere—restricts SETα expression at pre-blastocyst stages (Kinisu et al., 2021). Interestingly, a study has shown that SET prevents acetylation of KLF5 at its DNA-binding domain, negatively regulating its binding to DNA. A mutant version of KLF5 that cannot be acetylated does not depend on SET levels for transcriptional activation and cell growth (Miyamoto et al., 2003). This suggests that a positive feedback loop may exist, where increased levels of KLF5 reduce levels of SET, leading to increased activity of KLF5, thereby further reducing levels of SET.

In addition, we propose that downregulation of KLF5 after the 4-cell state allows full SETα expression where it can contribute to PrE specification. Finally, upon secure PrE specification, downregulation of FGF4 by rising levels of SETβ, along with KLF5 in the late blastocyst, may help to maintain epiblast identity (Fig. 6C-D). In future studies, performing scRNA-seq experiments following retinoic acid differentiation could highlight deficiencies in primitive endoderm formation in *Set*-KO cells. Likewise, attempting to generate XEN cells following rescue with SETα or SETβ could inform us further about the role of SET in primitive endoderm specification.

Our findings here shed light on differential regulatory roles of SETα/SETβ. The increased intrinsic disorder of the SETα N-terminus may allow for more promiscuous binding to promoters of stemness genes and their activation. Interestingly, this may explain why patients with elevated SETα relative to SETβ expression levels have poorer clinical outcomes in chronic lymphocytic leukaemia (Brander et al., 2019). The multifunctional nature of SET warrants further investigations of SET isoform-specific roles in different developmental and physiological contexts.

## Acknowledgements

This work was supported by the European Union’s Horizon Europe Research and Innovation Programme under the EIC Pathfinder Open grant agreement #101099654 (*RT-SuperES* to E.M.) and the Israel Ministry of Science (MOST)-DKFZ binational program #0005358 to E.M.). E.M. is the incumbent of the Arthur Gutterman Professor Chair for Stem Cell Research. P.S.L.L. was supported by the Marie Curie *EpiSyStem* ITN network (REP-PT-765966).

## Author contributions

P.S.L.L. performed all experiments and analyzed the data; P.S.L.L. and E.M. conceived the study and designed the analysis. P.S.L.L and E.M. wrote the manuscript. E.M. supervised the study.

## Data availability

The RNA-seq (Accession Number: GSE260530) and ChIP-seq (Accession Number: GSE260525) data generated for this study have been deposited at Gene Expression Omnibus (GEO).

## Supplementary Figure legends

**Supplementary Figure 1.**
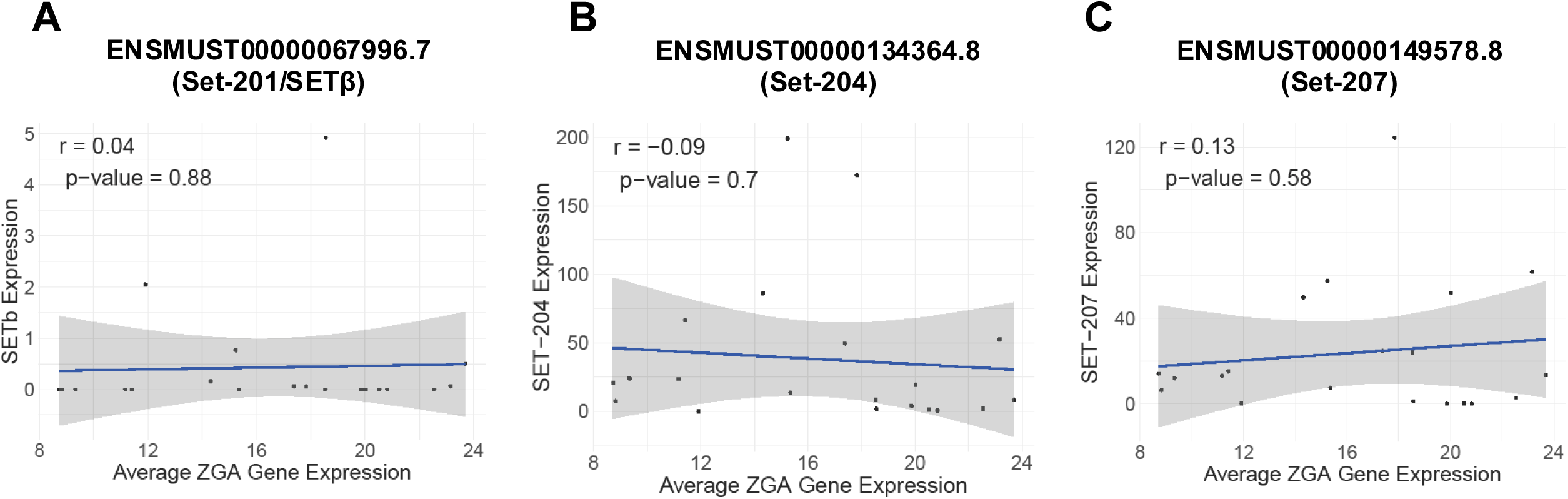
A) Correlation between SETβ transcript expression (TPM) and average ZGA expression (TPM). B) Same as (A) for protein coding SET transcript, Set-204. C) Same as (A) for protein-coding SET transcript, Set-207.

**Supplementary Figure 2.**
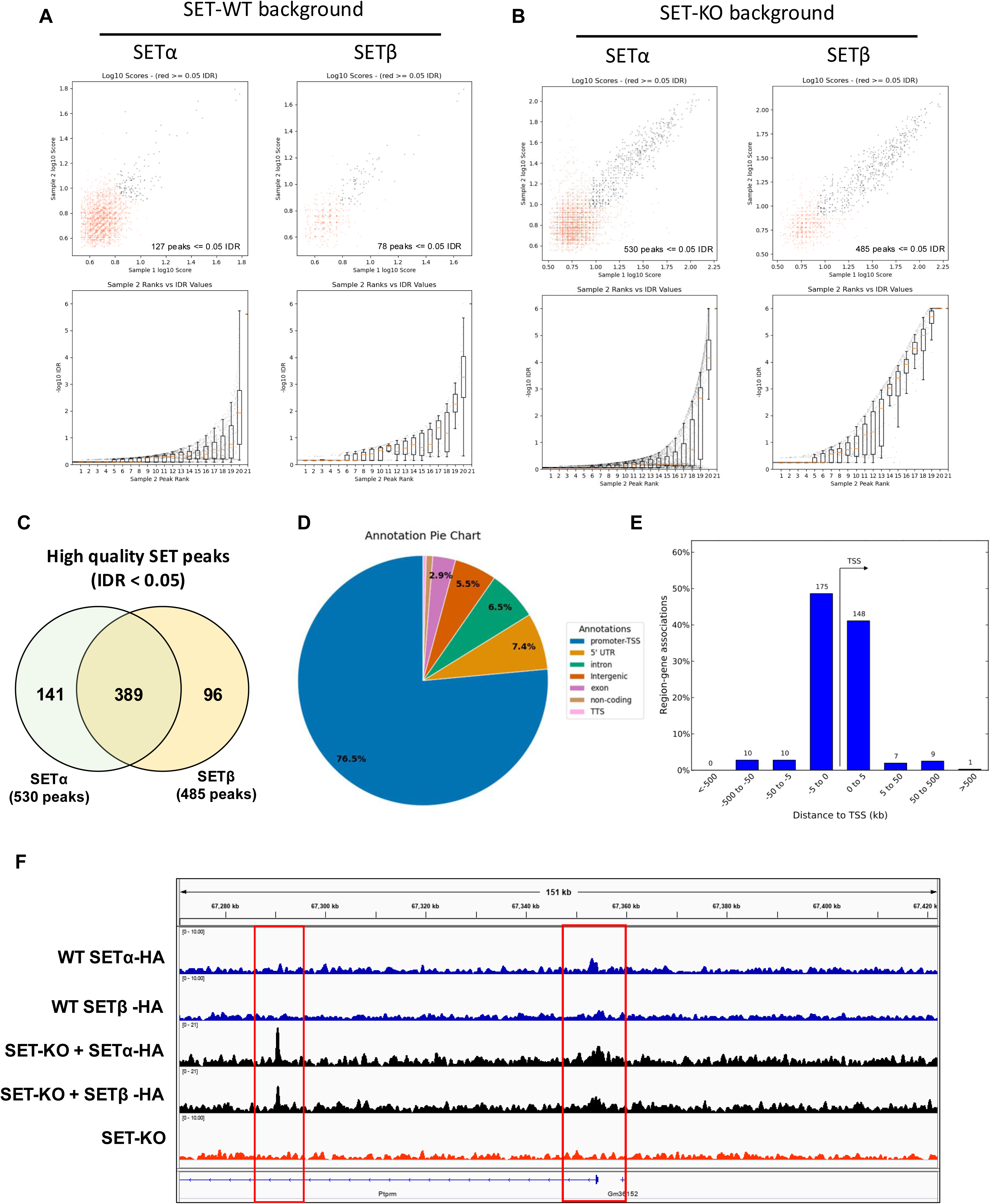
ChIP-seq for SETα and SETβ in WT and SET-KO background conditions. A) Reproducibility of peaks in two biological replicates for SETα (left) and SETβ (right) overexpression in SET-WT background using the IDR method. B) Reproducibility of peaks in two biological replicates for SETα (left) and SETβ (right) addback in SET-KO background using the IDR method. C) SET regions enriched in SETα and SETβ based on MAnorm-enrichment. D) Piechart showing proportion of peaks in different genomic features for SETα addback in SET-KO background. E) Location of SETα peaks relative to TSS for SETα addback in SET-KO background. F) IGV tracks comparing peaks from SET-WT and SET-KO ESCs with SETα/SETβ overexpression or SETα/SETβ addback, respectively.

**Supplementary Figure 3.**
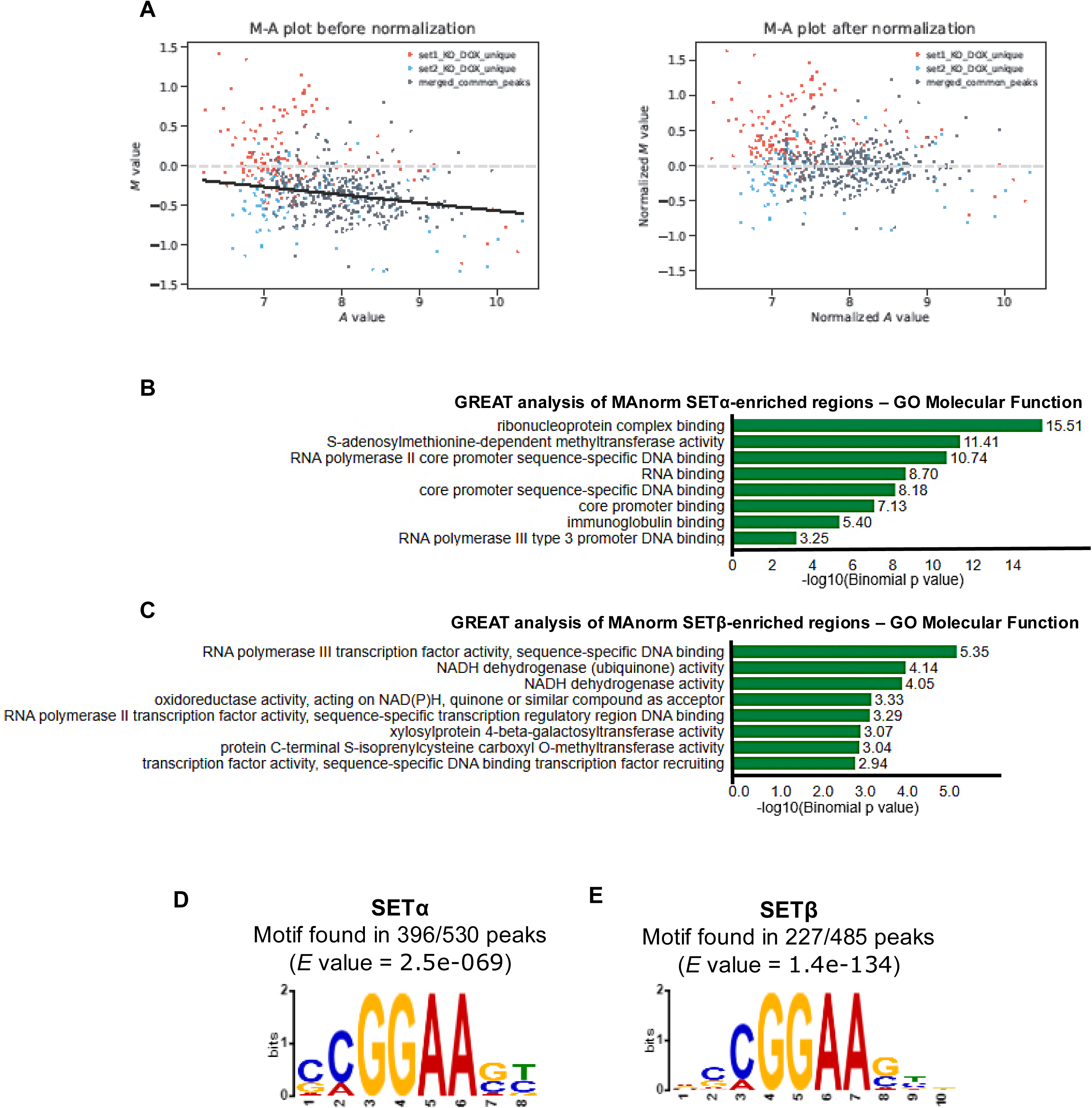
MAnorm normalization to identify enriched SET-isoform specific regions. A) MAnorm calculates normalization based on overlapping peaks (left). This is used to normalize M values (counts at each peak) to reveal enrichment of peaks specific to one sample (SETα or SETβ) (right). B) GREAT analysis of SETα-enriched regions for “GO Molecular Function”. C) Same as (B) for SETβ-enriched regions. D) Predicted binding motif for SETα peaks as identified by de novo motif discovery analysis using the motif-based sequence analysis tool MEME-ChIP. E) Same as (D) for SETβ peaks.

**Supplementary Figure 4.**
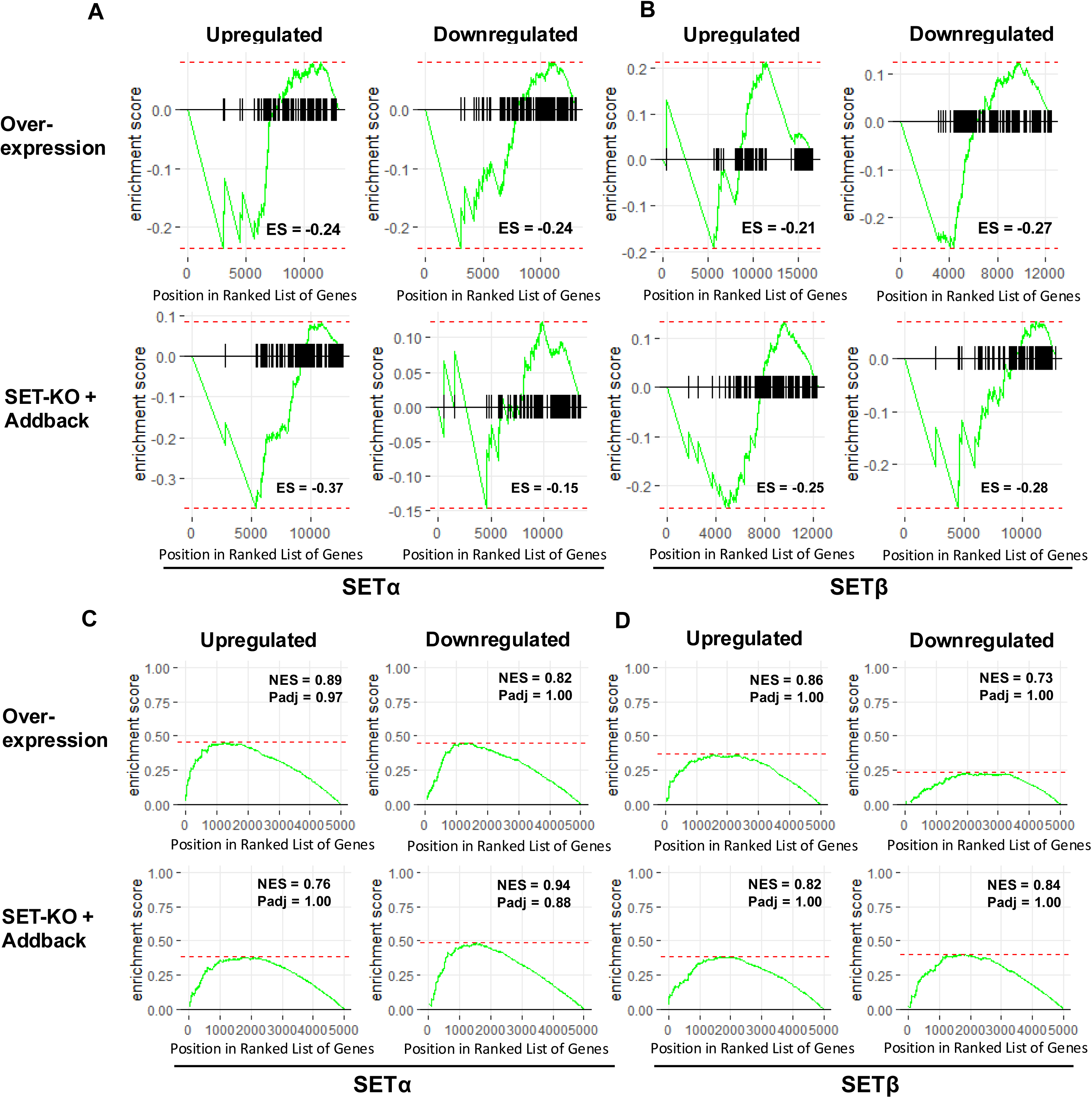
**Enrichment of ETS target genes in varying SET conditions.**A) GSEA plots displaying enrichment of ETS family target genes based on upregulated (left) or downregulated (right) genes from SETα overexpression (top) and SET-KO with SETα addback (bottom). B) Same as (A) for SETβ overexpression (top) and SET-KO with SETβ addback (bottom). C) GSEA plots displaying enrichment of ELK4 target genes based on upregulated (left) or downregulated (right) genes from SETα overexpression (top) and SET-KO with SETα addback (bottom). D) Same as (C) for SETβ overexpression (top) and SET-KO with SETβ addback (bottom).

**Supplementary Figure 5.**
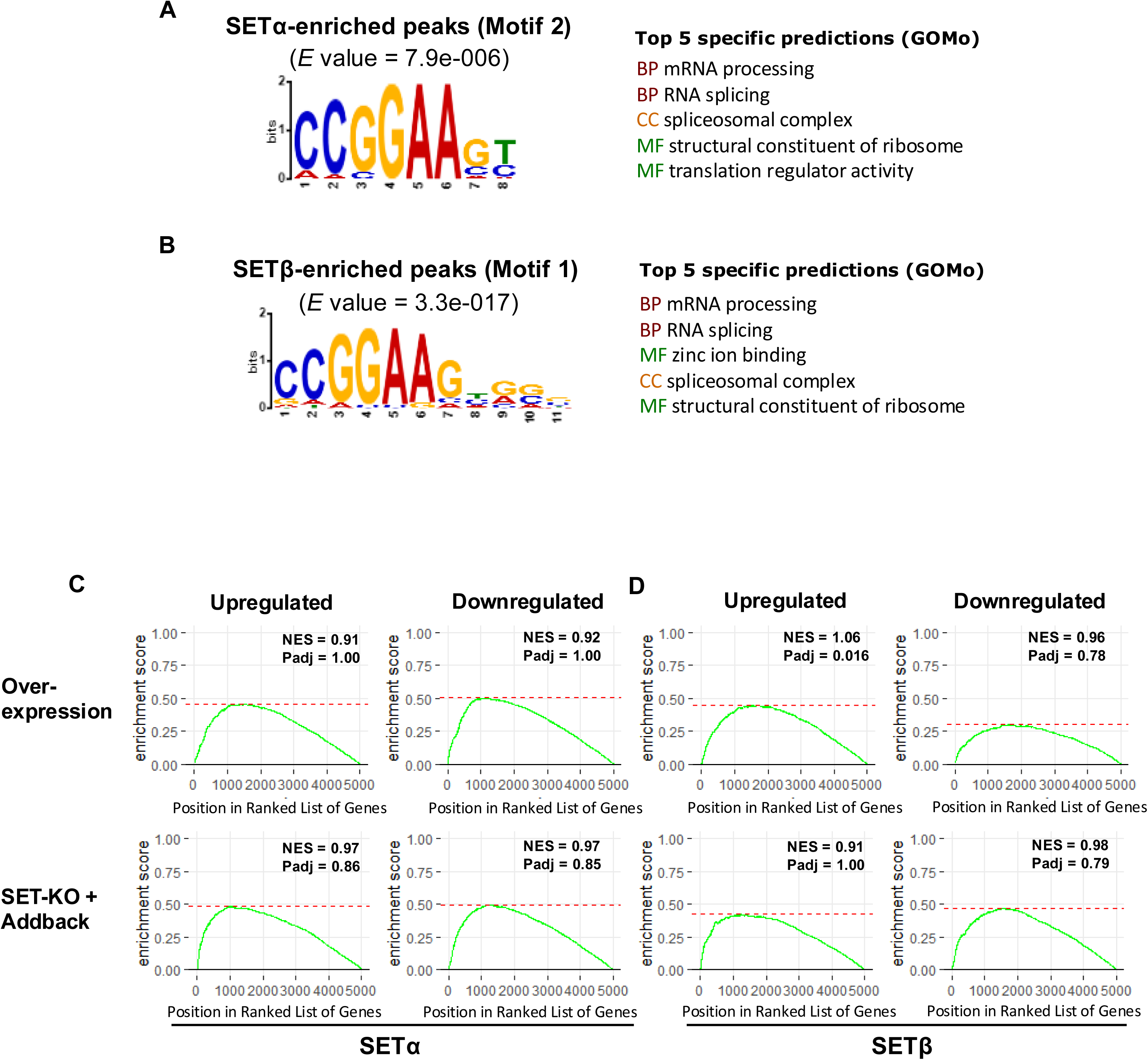
A) Second highest significant MEME-ChIP predicted de novo motif for MAnorm-enriched SETα regions and associated GOMo predicted regulatory gene ontology terms. B) Same as (A) for the highest significance motif for MAnorm-enriched SETβ regions. C) GSEA plots displaying enrichment of SETα-specific motif-containing genes (obtained with GOMo) based on upregulated (left) or downregulated (right) genes from SETα overexpression (top) and SET-KO with SETα addback (bottom). D) Same as (C) for SETβ overexpression (top) and SET-KO with SETβ addback (bottom).

**Supplementary Figure 6.**
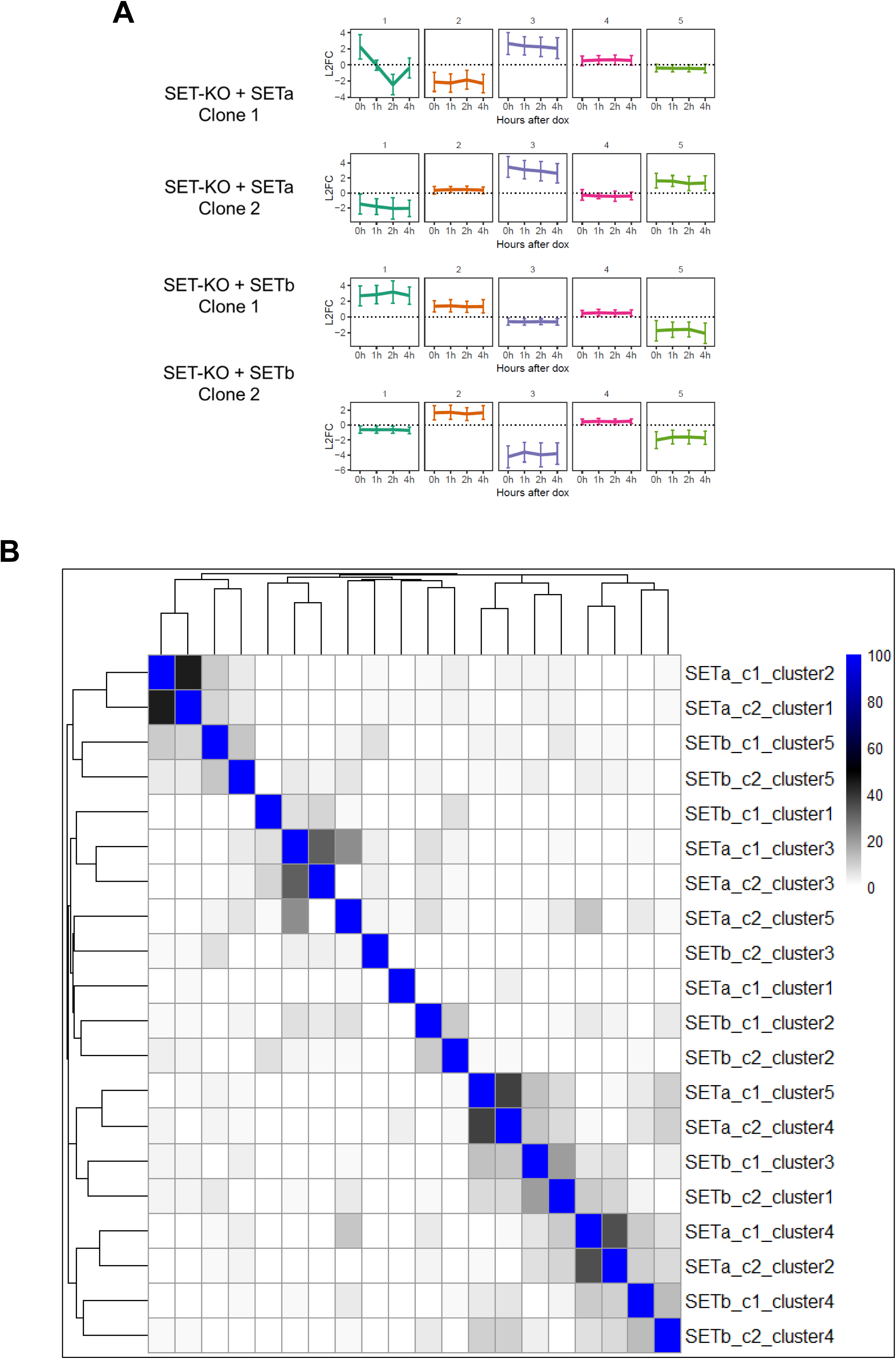
Gene expression patterns following time-course SETα/SETβ addback. A) K-means clustering of 5 clusters of genes with the same pattern of expression following SETα or SETβ addback in two independently derived clones. B) Heatmap with dendrogram displaying clusters that share genes in two independently derived clones.

**Supplementary Figure 7.**
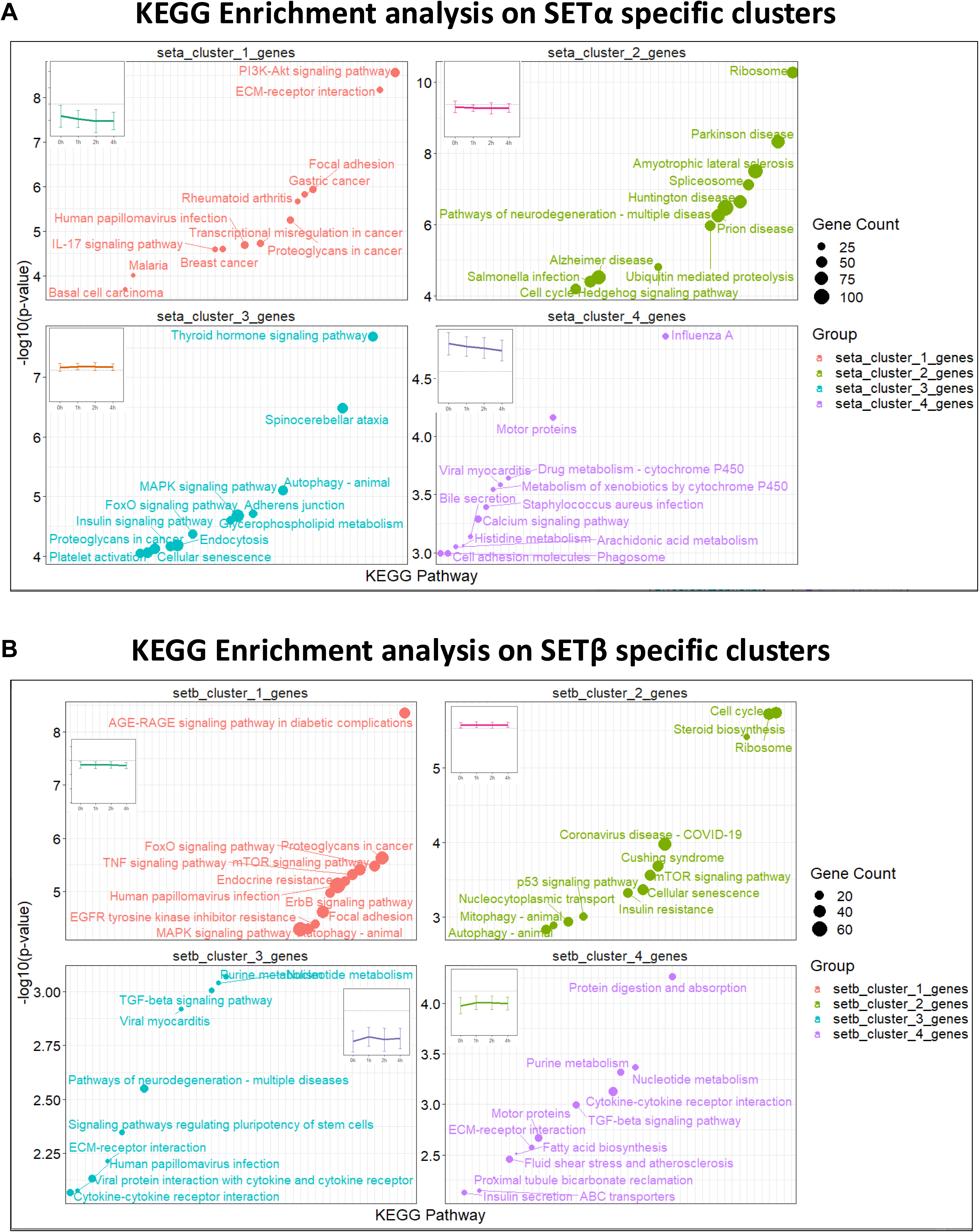
KEGG enrichment analysis on expression clusters following time-course SETα/SETβ addback. A) KEGG enrichment analysis for top 4 reprodicible clusters for SETα. B) KEGG enrichment analysis for top 4 reproducible clusters for SETβ.

**Supplementary Figure 8.**
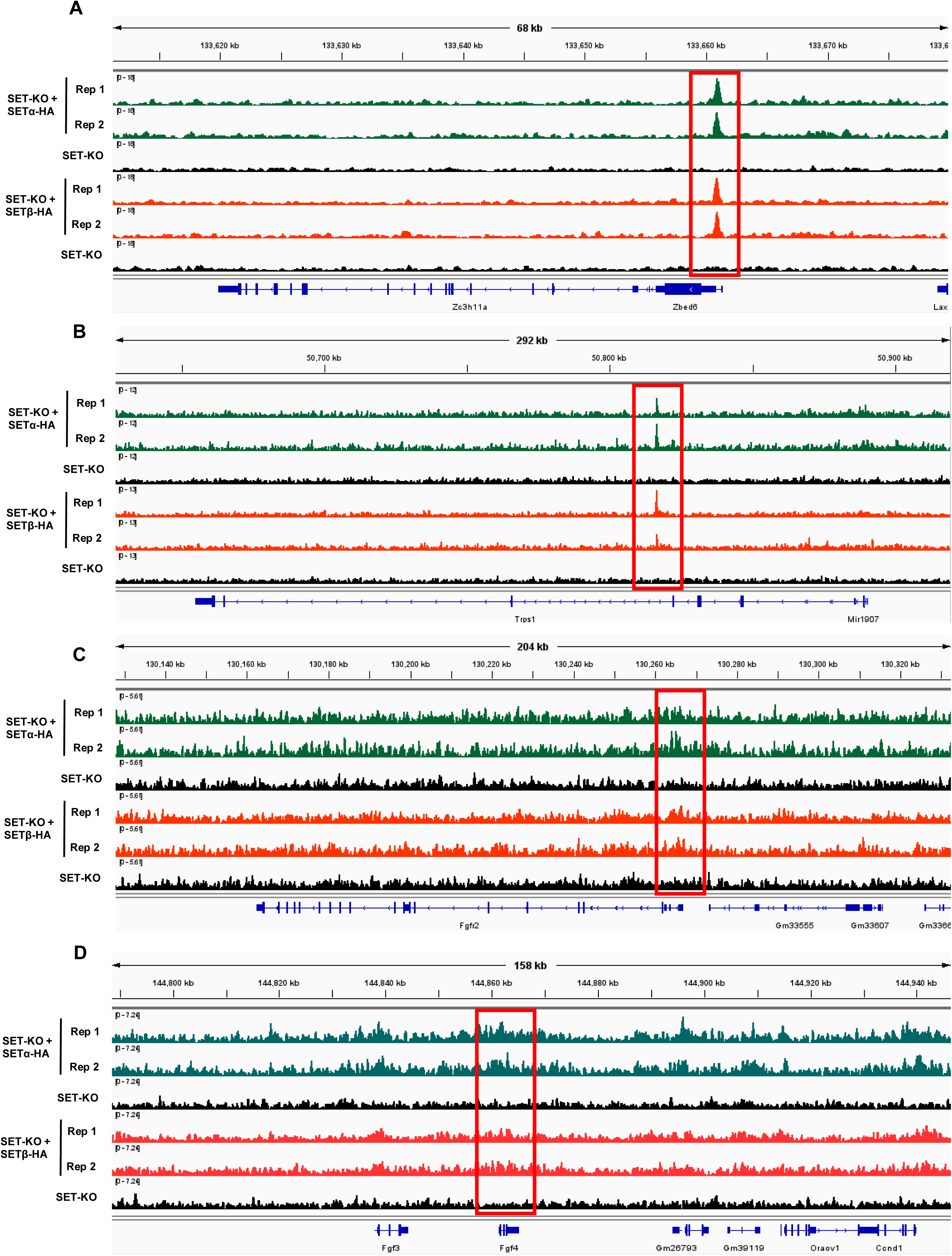
IGV tracks showing SETα enriched target genes. A) SET binding at the *Zc3h11a* gene locus. Two biological replicates are shown for each treatment group compared with one biological replicate for the respective background SET-KO cell line without Dox-induced SETα or SETβ expression. B) Same as (A) for *Trps1*. C) Same as (A) for *Fgfr2*. D) Same as (A) for *Fgf4*.

**Supplementary Figure 9.**
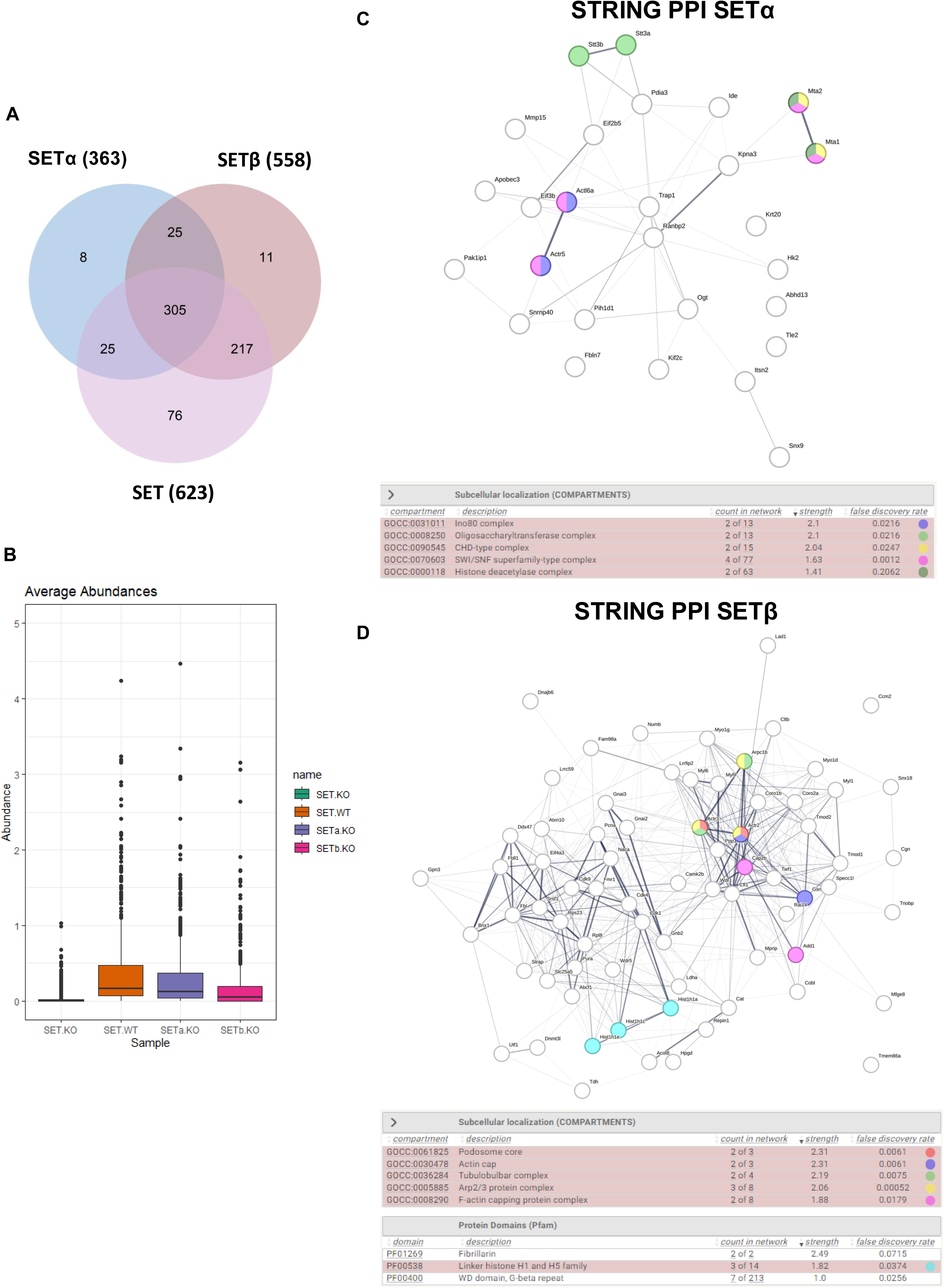
Analysis of isoform-specific interacting proteins from liquid chromatography coupled to tandem mass spectrometry experiments. A) Venn diagram showing overlap of proteins identified as peptide hits from mass spectrometry experiments. Peptide was counted if average emPAI – average emPAI of SET-KO experiment > 0. B) Abundance (peptide hits) of protein from liquid chromatography coupled to tandem mass spectrometry experiments for SET immunoprecipitation (IP) in different cell lines. C) Protein-Protein Interaction Networks Functional Enrichment Analysis (STRING) for SETα-interacting proteins (SETβ-KO). D) Same as (C) for SETβ-interacting proteins (SETα-KO).

